# Evolutionary Integration of a Foreign Aromatic Catabolic Pathway Drives Metabolic Trade-offs in *Methylobacterium extorquens* PA1

**DOI:** 10.64898/2026.03.02.709203

**Authors:** Alexander B. Alleman, Akorede L Seriki, Dawson J. Mathes, Monica J. Pedroni, Nkrumah A. Grant, Christopher J. Marx

## Abstract

Bacteria often acquire novel metabolic functions through horizontal gene transfer, allowing them to utilize new carbon sources. However, to benefit from these new pathways, they must be integrated with the host’s native metabolism. In nature, this process is fine-tuned via selection, enabling bacteria to exploit new niches. Alternate routes for pathway integration might yield distinct patterns of trade-offs, leading to differentiation within the adapting population. Understanding the metabolic mechanisms underlying these trade-offs provides insight into what maintains metabolic diversity in the environment. Lignin, a complex aromatic biopolymer, serves as an ideal substrate for exploring these questions, as monomers require complex metabolic pathways to break aromatic rings and be fed directly into central metabolism. In this study, we used the phyllosphere bacterium *Methylobacterium extorquens* PA1 to examine how a previously engineered catabolic gene cluster enabling lignin monomer utilization integrates with central metabolism. To this end, we experimentally evolved strains on lignin monomers vanillate (VA) and protocatechuate (PCA). Whole-genome sequencing revealed substrate-specific mutations that collectively reprogram stress responses and carbon storage regulation. These mutations resulted in myriad metabolic trade-offs: VA adapted strains showed diminished growth on PCA, PCA adapted strains largely cross-adapted to VA, and both VA and PCA evolution decreased growth rate on non-aromatic native substrates like succinate. These findings illuminate how evolution optimizes catabolic flux and resource allocation to efficiently integrate a foreign pathway following horizontal gene transfer, and the pleiotropic effects of such optimization.

**Importance:** Microbial use of aromatics derived from lignin, whether in natural ecosystems or in biomass conversion, has been a major research focus. While aromatic pathways are well-known to have broad substrate specificity, pulling in aromatic molecules at various levels of conversion, it is unclear how much evolutionary constraint there is upon adaptation to each of these steps. By experimentally evolving *Methylobacterium extorquens* carrying a foreign lignin-derived aromatic catabolic pathway, we demonstrate that adaptation involves not just increased pathway flux but also regulatory rewiring and reallocation of resources, especially through PHB metabolism and the TCA cycle. The results reveal substrate-specific trade-offs and cross-adaptation, illustrating how new functions reshape fitness landscapes. These insights on metabolic innovation, pleiotropy, and diversification following horizontal gene transfer have broad implications for microbial evolution and metabolic engineering.

## Introduction

Large amounts of the world’s available carbon are contained in molecules that are toxic and/or difficult to access metabolically. Lignin, a key component of plant cell walls and the second most abundant biopolymer on Earth, is notoriously recalcitrant, requiring specialized metabolic pathways for its degradation (1). Once broken down to its aromatic monomers, the carbon-rich structure and natural abundance make it an attractive carbon source for soil microbes (2, 3). Lignin is mostly composed of methoxylated aromatic monomers, which pose an additional problem since *O*-demethylation can produce the toxic byproduct formaldehyde. Organisms have evolved several strategies to mitigate the toxicity of formaldehyde. In model lignin-degrading organisms, such as *Rhodococcus jostii* RHA1 and *Pseudomonas putida* KT2440, growth on methoxylated aromatics, such as vanillate (VA), leads to the excretion of formaldehyde into the supernatant (4, 5). Laboratory evolution of the model *P. putida* KT2440 utilizing VA as its substrate revealed parallel mutations that enhanced the native glutathione-dependent formaldehyde detoxification pathway, demonstrating that formaldehyde toxicity is a bottleneck in methoxylated aromatic bioconversion (6).

Recently, *Methylobacterium*, a genus found predominantly in the phylloshpere and soil, has been discovered to utilize methoxylated aromatic compounds (4, 7). These *Methylobacterium* strains, which are natively methylotrophic, exhibit an enhanced ability to cope with formaldehyde produced from methoxylated aromatics. This is due to their native formaldehyde assimilation pathways and stress response genes, which enable growth on VA without releasing formaldehyde as a byproduct (4, 5, 8, 9). These aromatic-degrading *Methylobacterium* strains catabolize methoxylated aromatics through a unique β-ketoadipate pathway, which is encoded by a combination of the traditional *vanAB* and *pcaGHB* genes, the addition of *pcaL*, and a new gene, *kce*, replacing *pcaIJF* activity (**Figure 1A**). The gene *kce* is predicted to encode a β-ketoacid cleavage enzyme producing succinyl-CoA and acetoacetate from β-ketoadipate and acetyl-CoA (10). The production of succinyl-CoA, acetoacetate, and formaldehyde from VA catabolism integrates into the complex central metabolism of *Methylobacterium* using the serine cycle, the tricarboxylic acid (TCA) cycle, as well as the poly-3-hydroxybutyrate (PHB) and the ethylmalonyl-CoA (EMC) cycle (**Figure 1A**). Recently, work in *M. extorquens* SLI 505 demonstrated that the formaldehyde produced from the VanAB reaction is entirely dissimilated to CO_2_, allowing the carbon from the β-ketoadipate to be assimilated within central metabolism (9).

**Figure 1.**
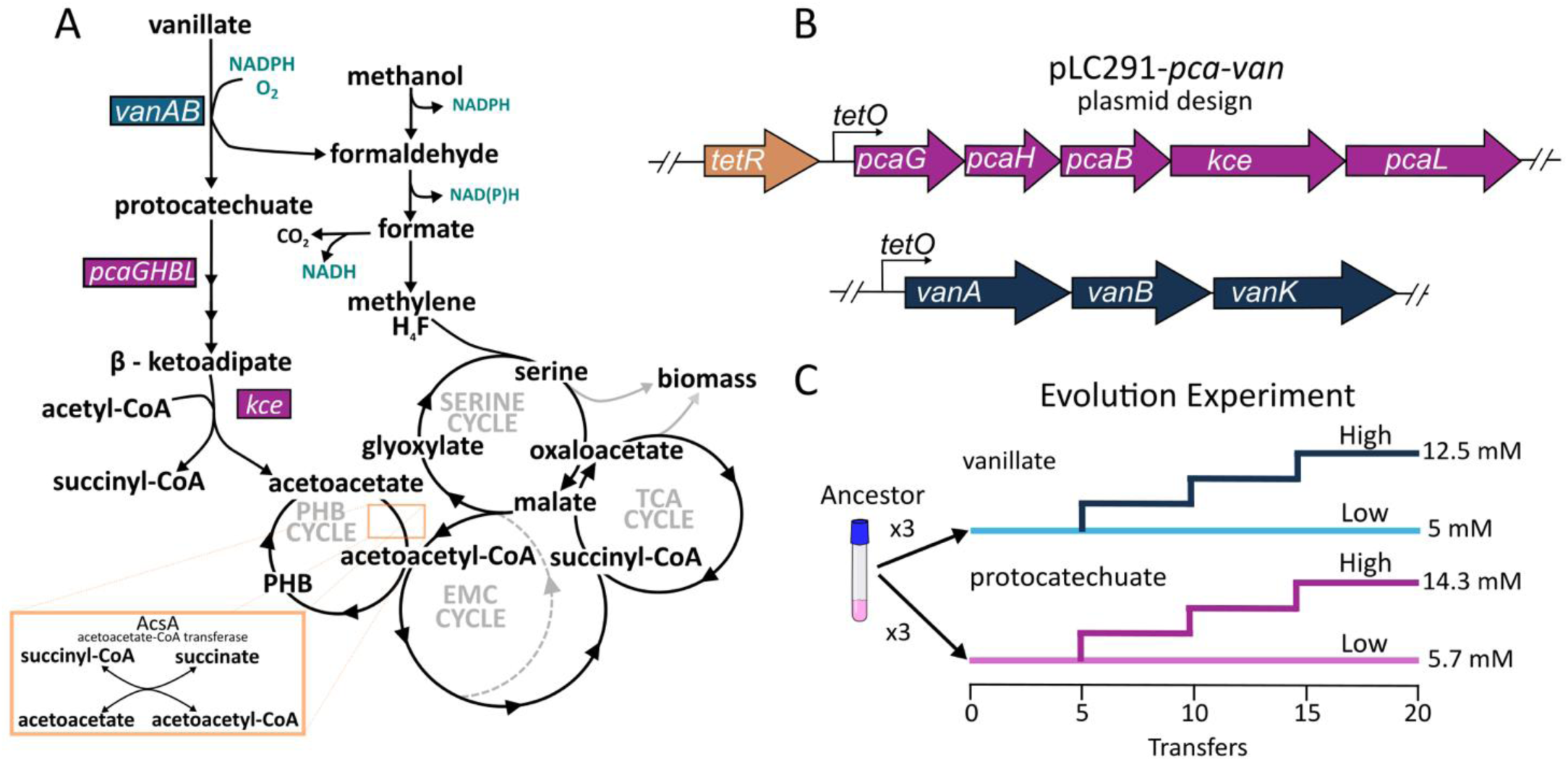
**A Central metabolic pathways of *M. extorquens* PA1.** Native pathways, including methanol metabolism, where C_1_ intermediates can be fully oxidized to CO_2_ or incorporated into biomass through the serine cycle and TCA cycle. When grown on C_1_-substrates, glyoxylate regeneration occurs through the ethylmalonyl-CoA (EMC) cycle, which requires the first two enzymes of the poly-3-hydroxybutyrate (PHB) cycle. The introduced pathway for aromatic assimilation (left) ends in the breakdown of aromatics VA and PCA to succinyl-CoA and acetoacetate through the novel enzyme Kce. Acetoacetate and succinyl-CoA swap carrier groups through the enzyme AcsA, which produces the final products acetoacetyl-CoA and succinate (insert box). **B) Plasmid design for aromatic gene engineering.** Both the *pca* and *van* operons from *M. nodulans* were placed on a TetR-based expression plasmid with two repression sites. **C) Evolution experiments**. Three replicates of the ancestor strain (*M. extorquens* PA1 with pLC291-*pca-van*) were serially transferred on either VA or PCA for laboratory evolution. After 5 transfers, each population was split into two; one with the same low substrate concentration, and one with a higher. The low population maintained a normalized carbon concentration of 40 mM (5 mM VA, 5.7 mM PCA). At the same time, the high population experienced an increase in substrate concentration at every 5 transfers, reaching a final normalized carbon concentration of 100 mM (12.5 mM VA, 14.3 mM PCA). A total of 20 transfers were made, equivalent to 120 generations.

Previously, our lab reported that genes for VA catabolism from *Methylobacterium nodulans* ORS 2060 can be successfully introduced into the laboratory model *M. extorquens* PA1 (5). The genetic engineering preformed was analogous to natural examples of recent acquisition of aromatic utilization via horizontal gene transfer (HGT) by other *Methylobacterium* strains (4, 7). The introduced plasmid, pLC291-*pca-van* has two distinct regions of the *M. nodulans* chromosome cloned co-directionally, but each with its own tetracycline transcriptional promoter (*P_R/tetO_*) (**Figure 1B**). The TetR repressor present thus controls the expression of both operons separately, which can be regulated by anhydrous tetracycline (aTc) (11). Upon introduction of pLC291-*pca-van* plasmid, the engineered *M. extorquens* grew as fast as *M. nodulans* on elevated concentrations of VA without excreting formaldehyde. The ability of *M. extorquens* PA1 to utilize the pathway without major adaptation was surprising, but it allows us to ask how a foreign pathway with high flux can integrate into native metabolic networks after an HGT event.

A central challenge for organisms evolving in environments with multiple substrates is balancing specialization and generalization, as mutations that improve the utilization of one compound may exert antagonistic pleiotropic effects on others (12–14). Tradeoffs such as this are frequently observed, such as *M. extorquens* AM1 strains, which evolved on succinate, often lost their ability to consume methanol (15) because mutations that disabled C_1_ assimilation were beneficial during growth on succinate (16). Alternatively, there may be instances of synergistic pleiotropy, where adaptation to one substrate yields gains on others. For example, the evolution of *M. extorquens* AM1 with a foreign formaldehyde oxidation pathway on methanol led to similarly large gains on an alternate substrate, methylamine (17), because of the strong overlap in the pathways for utilizing both substrates. However, we lack a similar understanding of the pleiotropic effects when populations acquire new pathways and evolve to better utilize novel substrates.

In this work, we evolved *M. extorquens* PA1 containing a novel aromatic catabolic pathway on either VA or PCA. Adaptation to foreign pathways can lead to sets of unpredictable pleiotropic effects. First, we hypothesized that because VA and PCA catabolic pathways overlap except for the additional toxicity of the demethoxylation step, evolution on VA would lead to increased fitness on PCA, whereas the reciprocal condition would lead to antagonistic pleiotropy. Second, since both aromatics enter the central metabolism via the PHB, EMC, and TCA cycle, we expect there could be pleiotropic effects on native central metabolites, such as methanol and succinate. Overall, we found that evolution to VA and PCA led to distinct patterns of adaptation and a mixed set of both antagonistic and synergistic effects on alternate substrates. These adaptations were driven by recurrent mutations in key regulatory and metabolic genes related to aromatic breakdown, stress response, and carbon storage. Our research highlights how selection modulates gene expression and physiology to better utilize novel metabolic pathways and reallocate resources. This differentiation during evolution and antagonistic pleiotropy can help explain the creation of different niches within a rapidly evolving population.

## Methods

### Bacterial strains, media, and chemicals

The *M. extorquens* PA1 strains used in our study, including the ancestors and evolved progeny, are stored frozen at −80 °C with dimethyl sulfoxide (DMSO) added as a cryoprotectant (final concentration 8% v/v). We revived strains by inoculating 10 µl of bacterial stock into 5 mL of minimal MPIPES media (18) with 3.5 mM succinate and 50 µg/mL kanamycin. All strains used in this study are listed in **Table S1**. We incubated the resulting cultures at a 45° angle in a shaking incubator set to 250 RPM and 30 °C (henceforth, standard conditions), where they reached the stationary phase within 48 hours. To precondition the strains for the growth conditions of the evolution experiment, we diluted the cultures 1:64 in 5 mL of MPIPES medium containing 5 nM aTc, along with the aromatic compounds PCA or VA, both normalized to 40 mM carbon (5 mM VA and 5.7 mM PCA). The addition of kanamycin was required for growth on succinate, whereas growth tests on VA or PCA did not require it, as the plasmid was selected solely to maintain substrate use.

The plasmid, pLC291-*pca-van*, was introduced into CM4916 (CM2730 with Δ*phaC*) via triparental conjugation using the helper plasmid pRK2073 (19). The resulting transconjugants were screened on selection plates containing MPIPES, succinate, and 50 μg/mL kanamycin, and verified by Sanger sequencing, generating strain CM4672.

### Experimental evolution

Three clonal replicate populations (A to C) of CM4669 were initiated in MPIPES supplemented with either 5 mM VA or 5.7 mM PCA (each 40 mM C) plus 5 nM aTc to provide modest induction of the *van* and *pca* operons. The culture volume was 5 mL, maintained in 18×150 mm borosilicate glass culture tubes, and incubated under standard conditions. We serially propagated cultures by transferring a 1/64 dilution weekly, providing for six net generations per week. After the fifth transfer, each of these triplicate populations was split into two sub-populations. The ‘low’ populations stayed at 5 mM VA or 5.7 mM PCA for the rest of the transfers. While the ‘high’ populations for both VA and PCA experienced a concentration increase of 2.5 or 2.9 mM, respectively (20 mM increase in C atoms, **Figure 1C**). These populations experienced further concentration increases after the 10^th^ and 15^th^ transfers, respectively, until they reached the final concentrations maintained until the end of the evolution experiment (12.5 mM VA or 14.3 mM PCA).

### Growth quantification and measurement of specific growth rate

Optical density (OD) measurements were recorded in a 96-well plate using a Synergy H1 Plate reader (BioTek), at 600 nm (OD_600_), using a protocol previously described (5). Raw OD values were processed using R (version 2023.06.1+524) with the growthrates, tidyverse, and ggplot2 packages, also using a previously described method (5). Blank wells, containing only media and carbon source without cells, were used to correct for background absorbance by subtracting the OD_600_ value from all sample readings.

### Genome sequencing analysis

Genomic DNA from evolved isolates of the engineered *M. extorquens* was extracted using the MasterPure complete DNA extraction kit (Biosearch Technologies, Shanghai, CN). Libraries were prepared for Illumina sequencing by fragmenting DNA to ∼350 bp via sonication, followed by end-polishing, A-tailing, and ligation of Illumina-compatible adapters. Libraries were bead-cleaned to remove short fragments, and quality was verified using fluorometric analysis (Qubit 2.0, Thermo Scientific) and fragment analysis. Quantification of sequenceable libraries was performed via qPCR.

Sequencing was conducted on a full 2×300 bp Illumina Nextseq run (SeqCoast Genomics, NH, USA). Reads were demultiplexed initially with bcl2fastq2. Reads were aligned to the *M. extorquens* PA1 reference genome (Assembly: GCF_000018845.1) and the plasmid pLC291-*pca-van* using BreSeq for variant calling and mutation analysis (20). All genomes are available at the SRA BioProject PRJNA1284354.

### RNA isolation and real-time reverse transcription-quantitative PCR (RT-qPCR) analysis

Total RNA was extracted from 5 mL of exponentially growing cells (defined as half-maximal optical density at 600 nm, OD_600_) using the RNeasy Mini Kit (QIAGEN), and residual genomic DNA was removed using DNase I. Purified RNA was then utilized for RT-qPCR, with the gene-specific primers (Supplementary Table S3). Briefly, 1000 ng RNA was reverse transcribed into cDNA using the SuperScript™ III First-Strand Synthesis SuperMix for qRT-PCR (ThermoFisher), according to the manufacturer’s instructions. Two-step real-time PCR was performed on a Quant Studio 3 system with the PowerTrack™ SYBR Green Master Mix for qPCR (ThermoFisher), according to the manufacturer’s instructions. The gene *rpsB*, encoding the 30S ribosomal protein S2, served as the reference gene (21). The average threshold cycle (Ct) value for each gene was calculated from triplicate reactions, using a previously described method (22). The ΔCt value represents the difference between the Ct of the target gene and the Ct of the reference *rpsB* gene, while the ΔΔCt value describes the difference between the ΔCt of the ancestor and that of the evolved strains. The difference in expression was calculated using the Log_2_^ΔΔCt^ method. The resulting data were analyzed by a t-test of three biological replicates. P < 0.05 was considered statistically significant.

### AlphaFold Details

To analyze the effects of mutations on protein structures, the AlphaFold Server was used, with the AlphaFold 3 model (23). For the porin structure, the amino acid sequence was taken from the reference genome, and the protein was modeled as a trimer. When interpreting the mutation used in our experiment, we used the top-ranking structure (pTM = 0.88) for analysis.

### PHB quantification

Cells were grown in a 30 mL volume in a 250 mL Erlenmeyer flask with the desired carbon source in MP media, shaking at 250 RPM at 30 °C till reaching the stationary phase or taken at exponential phase when cells reached a half-maximal OD. All 30 mL of cells were centrifuged (4k RPM for 10 min), then the pellets were stored at −80 °C. Cell pellets were then lyophilized until the cell dry weight no longer changed, and the pellet mass was recorded. For PHB extraction, we followed an alkaline extraction method to quantify the monomer crotonic acid (24, 25). Briefly, cell pellets were mixed with 0.5 mL H_2_O and 0.25 mL 2 N NaOH, vortexed, and then boiled for 1 hr. The mixture was then neutralized with 0.25 mL 2 N H_2_SO_4_, centrifuged to remove cell debris, and filtered through 0.2 µm filter prior to HPLC analysis. The HPLC method consists of injecting 10 µL of sample into a mobile phase containing 10% acetonitrile and 90% H2O with 0.025% H_2_SO_4_. The mobile phase was run at 1 mL/min for 10 min through a Luna C-18(2) column (Phenomenex, Torrance, CA, USA) with an oven temperature of 60 °C. The quantification of sample PHB was performed using a standard curve of known PHB polymer. PHB extraction was compared to known crotonic acid standards to determine extraction efficiency.

## Results

### Experimental evolution of engineered *M. extorquens* on VA or PCA improves growth in selective environments

To understand how *M. extorquens* adapts to the acquisition of a foreign aromatic catabolism pathway, we evolved an engineered strain with the pLC291-*pca-van* plasmid on either VA or PCA as the sole carbon source (**Figure 1C**). Three populations were established on a low concentration of either substrate. After 30 generations, each of these sub-populations were split into two: ‘low’ sub-populations that would maintain their concentration through 120 generations, and ‘high’ sub-populations that would increase by 20 mM total carbon every 30 generations until reaching the maximum evolved concentrations of 100 mM total carbon (12.5 mM VA or 14.3 mM PCA). Isolates from both low- and high-subpopulation groups for each compound were isolated and tested for growth in their evolved environments relative to their common ancestors. Evolved isolates exhibited, on average, an improvement in growth rate of 75% (range: 38-115%) in their evolved substrate and concentration (**Figure 2, Table S4**).

**Figure 2.**
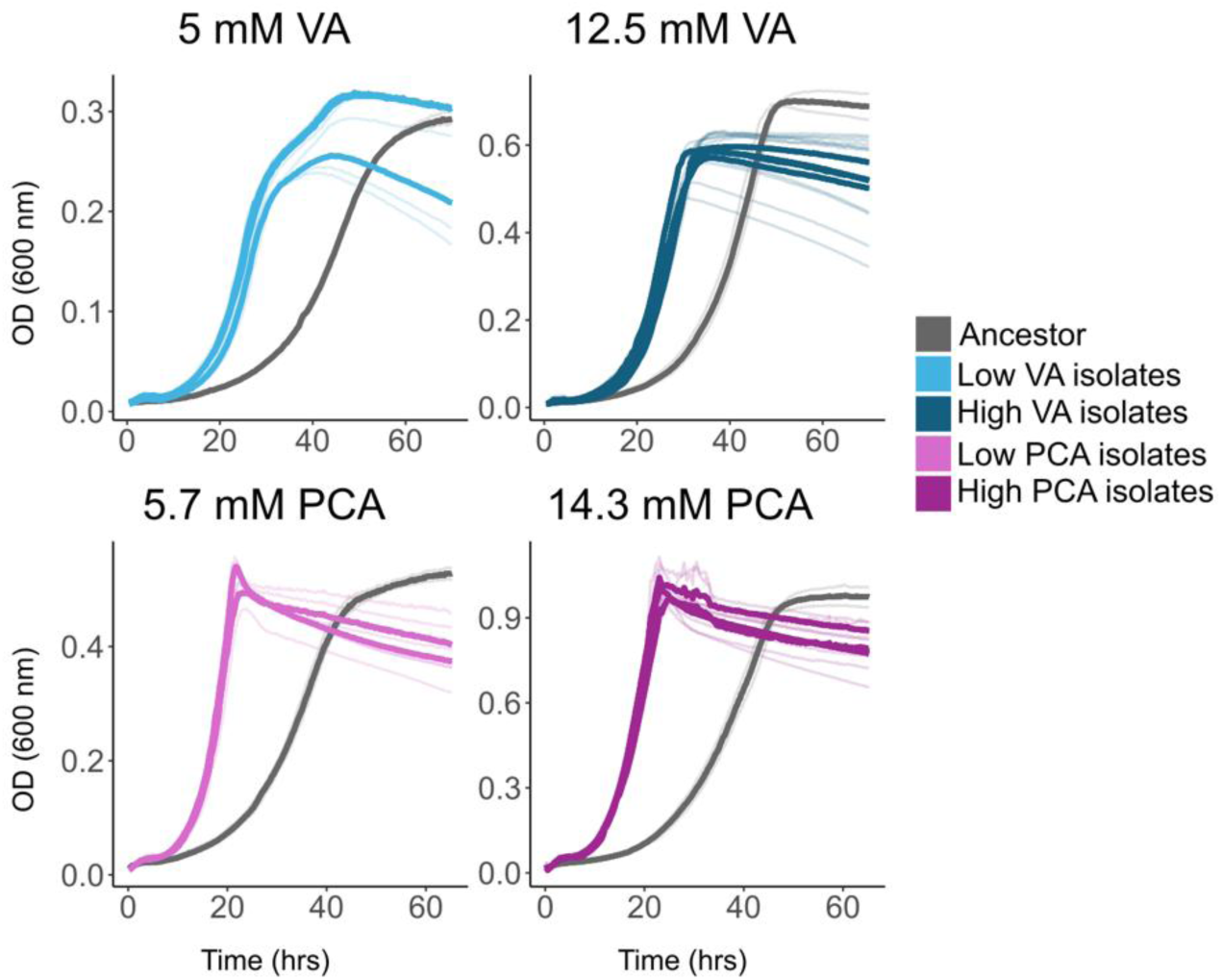
Evolution substantially improved growth in selective conditions. Growth curves of the ancestor strain (grey) compared to isolates taken from populations evolved in either VA (blue) or PCA (purple) to a low (light shade) or high (dark shade) concentration. Growth curves were performed in triplicate (light background), and the average OD was recorded (dark foreground).

To understand the genetic basis of adaptation, we performed clonal whole-genome sequencing for 3 isolates from each replicate population (A, B, C) of each of the 4 regimes (low and high, PCA and VA). Across replicate populations, we observed a high degree of parallel evolution, with recurrent mutations concentrated in a shared set of genes and regulatory elements. Mutations arose in both the plasmid encoding the heterologous *van* and *pca* operons and in the chromosome targeting genes associated with metabolic and stress adaptation (**Table 1, Table S3**). In the following text, we will first describe how plasmid-borne mutations affecting the TetR/*tetO* regulatory system influenced adaptation to VA and PCA, followed by an analysis of chromosomal mutations that contributed to the evolved phenotypes.

**Table 1.**
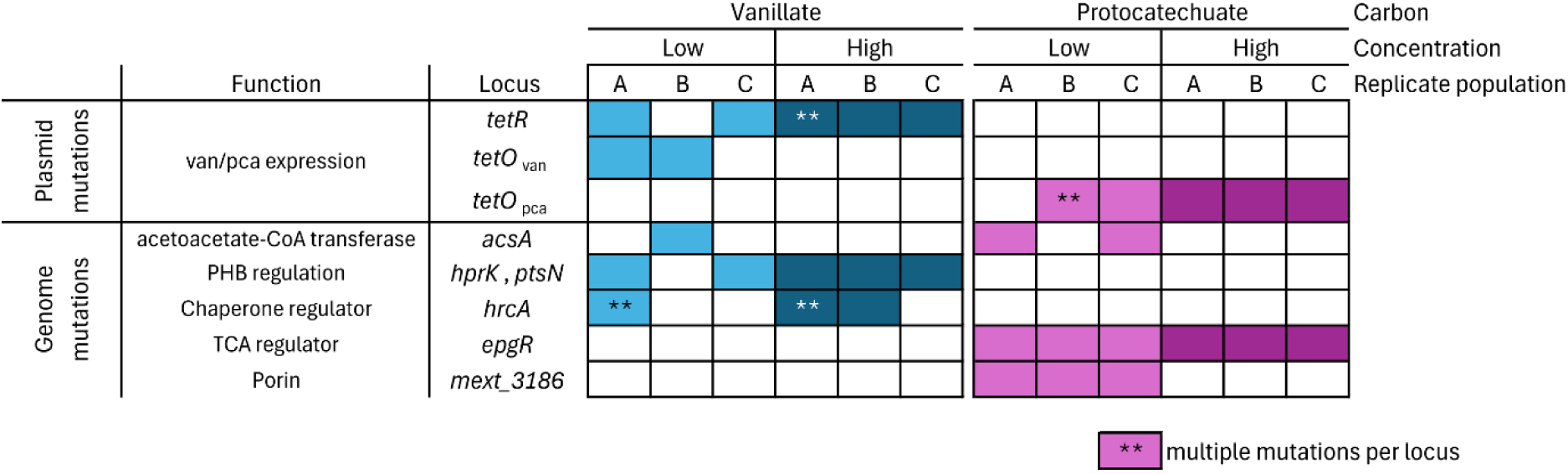
Mutations identified in evolved isolates. Three isolates from each population (A, B, C) and condition (VA, PCA) were sequenced and compared to the ancestor. Many loci occurred in parallel in different populations. Boxes are filled if at least one of the three isolates had a mutation at that locus, whether it is intragenic or in the predicted promoter region. We denote polymorphism for a single locus within a replicate population with a double asterisk. Color shading demarks low- or high-carbon conditions.

### Adaptive mutations in plasmid-borne TetR/*tetO* regulatory elements modulate the expression, which modulates the regulation of the foreign catabolic pathway

Nearly all evolved isolates acquired mutations in the TetR/*tetO* regulatory system, affecting either the repressor or one of the associated promoter regions (**Table 1**). These mutations represent the two ways to alter the expression levels of the newly introduced pathway. Mutations in the TetR repressor protein affect gene expression of both the *van* and *pca* clusters by altering the repressor’s binding to the tet operator (*tetO*) and/or the inducer molecule aTc’s binding to TetR. Mutations in *tetO* affect gene expression (by altering TetR repressor binding) only in the *van* or *pca* cluster, depending on whether the mutation occurs in the *tetO_van_* or *tetO_pca_* site. Mutations found in TetR repressor were only found in VA growth conditions and were nonsynonymous mutations in the protein coding region – H44Y, W76R, and R105W. The *P_R-tetO_* mutations occurred both at sites upstream of *vanABK* (*tetO_van_*) and upstream of *pcaHGBL-kce* (*tetO_pca_*); these mutations disrupted the palindromic sequence of the operator (**Figure 3A**).

**Figure 3.**
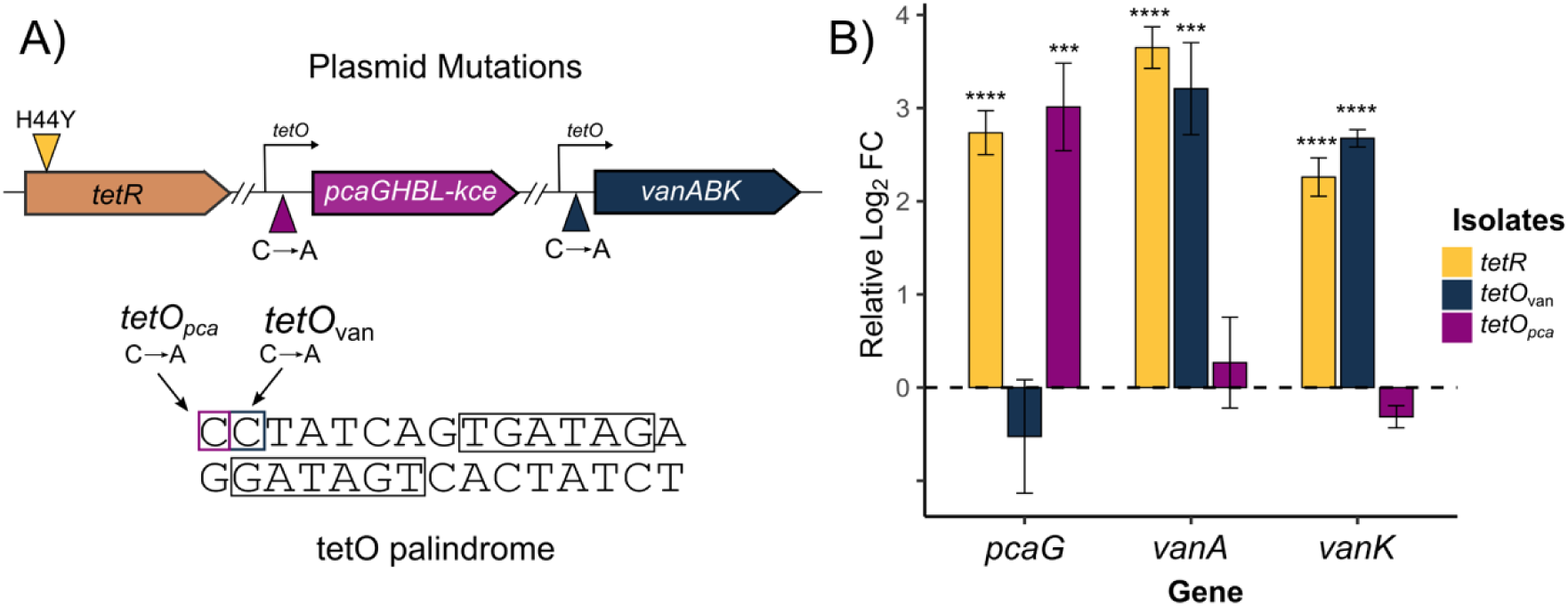
Mutations in the regulation of engineered genes affect expression. **A)** Plasmid organization and mutations present in evolved strains. A non-synonymous point mutation in *tetR* creates a H44Y change in a known residue required for hydrogen bonding to the *tetO* operator. The *tetO* mutations occur right at the start of the palindrome. **B)** Relative gene expression (log₂ fold change) of *pcaG, vanA*, and *vanK* in evolved isolates compared to the ancestor strain. Bars represent the mean relative log₂ fold change ± standard error (SE) from three biological replicates. Asterisks (*) indicate statistically significant differences in gene expression relative to the ancestor (*** p < 0.001, **** p < 0.0001, two-tailed t-test).

For both low and high PCA evolved isolates, mutations were only seen in the *tetO_pca_* sequence, which resulted in increased expression of the *pca* cluster (measured through *pcaG* expression) relative to the ancestor and had no impact on expression of the *van* genes (**Fig 3B**). However, in low- and high-Va evolved isolates, mutations were observed in both the VA utilization-specific *tetO_van_* site and the coding sequence of the TetR repressor protein. Mutations in the TetR repressor protein resulted in increased expression of both *van* and *pca* gene clusters, while mutations in the *tetO_van_* site resulted in increased expression of only the *van* genes (**Fig 3B**). The fact that TetR repressor mutations were specific to the VA conditions suggests both *pca* and *van* genes are selected for in VA conditions.

### Chromosomal mutations reflect regulatory, metabolic, and stress response adaptations

In addition to mutations arising on the plasmid, all isolates acquired one or two mutations on the chromosome. Only one chromosomal locus was targeted across both VA and PCA conditions (**Table 1**). This gene encodes the 3-oxoacid transferase/acetoacetate-CoA transferase named *ascA.* It is the last step of the PHB cycle, which transfers a CoA moiety from succinyl-CoA to acetoacetate, producing succinate and acetoacetyl-CoA (**Figure 1A**, inset) (26). The end of the β-ketoadipate pathway in *Methylobacterium* is predicted to generate acetoacetate and succinyl-CoA; this would also implicate AcsA in generating acetoacetyl-CoA and succinate by exchanging the CoA unit from succinyl-CoA (26) (**Figure 1A**, and **Figure 4A**). The exact same point mutation appeared independently in both VA and PCA evolved isolates and is located 120 bp upstream of the start codon, suggesting it alters the expression of *acsA*. The isolate with this upstream mutation showed an increased expression of *acsA* with a log_2_ fold change of 1.26 relative to ancestor (p < 0.001) (**Figure S1**), suggesting a potential link between aromatics metabolism and PHB accumulation that we explore further below.

**Figure 4.**
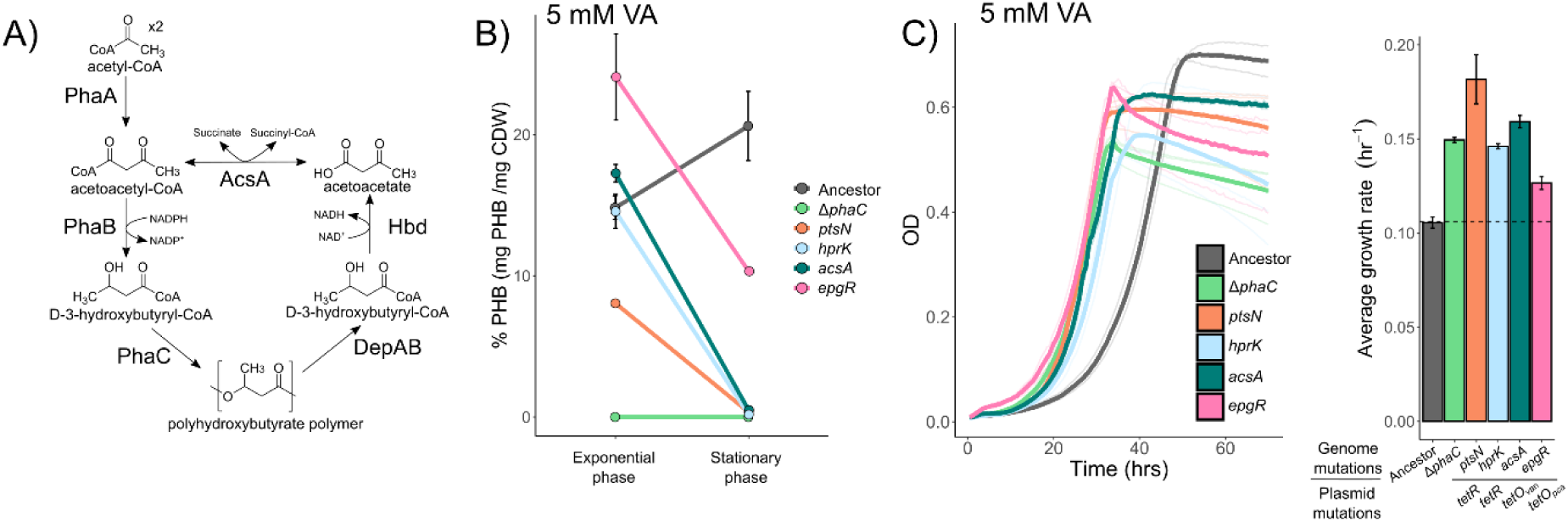
PHB accumulation affects growth on aromatics. **A)** The PHB cycle in detail. Carbon and electrons are stored in PHB through the condensation of two acetyl-CoA to make acetoacetyl-CoA. From there, this is polymerized to make PHB through PhaC. Depolymerization creates acetoacetate, where the final enzyme, AcsA, transfers a CoA from succinyl-CoA to acetoacetate, resulting in the formation of succinate and acetoacetyl-CoA. **B)** Total percent PHB of ancestor and isolate strains at exponential and stationary phase, both under evolved conditions of 5 mM VA. **C)** Growth curve of evolved isolates, the Δ*phaC* construct, and the ancestor at 5 mM VA, with each isolate colored by major genome mutation, and growth rates are also shown

One function that was uniquely targeted in VA-evolved populations was the PTS phosphorylation cascade, involved in Carbon:Nitrogen (C:N) ratio regulation and PHB mobilization in Hyphomicrobiales (27). These mutations were found in separate operons, with one, *Mext_1236* (*hprK*), containing the full gene set for the PTS phosphorylation cascade, complete with membrane proteins and other *pts* genes (**Figure S2A**). The other mutation in *Mext_2503* (*ptsN*) is located within a different operon that lacks a complete set of potential PTS phosphorylation cascade genes (**Figure S2B**). These signaling proteins have been implicated in PHB regulation in other alpha-proteobacteria, further suggesting that PHB cycling is relevant for aromatics use (27).

Another locus which was repeatedly targeted in VA-evolved populations was *hrcA* (*Mext_0399*), which is predicted to encode a heat-inducible transcription repressor. In related Hyphomicrobiales, the HrcA repressor binds the cis-acting element called CIRCE (for <u>c</u>ontrolling inverted repeat of <u>c</u>haperone <u>e</u>xpression) (28, 29). This CIRCE element is also present in the transcriptional start site upstream of the two *groES/EL* operons in *M. extorquens PA1* (**Supplemental material A**). Given the known role of HrcA in repressing *groE* chaperone, managing protein damage and misfolding (30), these mutations may alleviate general protein misfolding caused by VA or a specific challenge in folding VanABK. Expression data showed no significant change in the *groEL1* transcript compared to the ancestor despite having the Δ96 bp deletion within the *hrcA* gene (**Figure S1**).

### Evolution on PCA leads to parallel mutations in the novel gene *epgR*

For the PCA-evolved populations, an allele was identified in all three populations, encompassing both low- and high-concentration subpopulations, and was present in 15 of the 18 PCA-evolved isolates. This allele was *Mext_1642*, a gene encoding a TetR-family/AcrR-like transcriptional regulator (31). Since this gene is unique to the Methylobacteriaceae family **(Figure S2C)** and only shares partial homology with closely related *acrR* genes, we have decided to name this gene *epgR* for <u>e</u>nhanced <u>P</u>CA <u>g</u>rowth regulator. In total, three different mutations (−10, −11, or −67 bp) were selected for within the transcriptional binding sites upstream of the start codon of *epgR* (**Supplemental material B**). Each of these mutations enhanced growth on PCA to a similar extent (**Figure S3**). Interestingly, *epgR* is located directly up stream of the TCA cycle genes encoding malate dehydrogenase (*mdh*), oxoglutarate dehydrogenase (*sucAB*), and succinyl-CoA synthase (*sucCD*) (**Figure S2C**). While statistically insignificant, RT-qPCR showed a small decrease in expression of *sucAB* and *sucCD* relative to the ancestor (**Figure S1**).

Another mutational target common among the low PCA isolates was *Mext_3186*, a gene encoding a putative outer membrane porin. Three isolates from independent populations contained small in-frame deletions (Δ6, Δ6, or Δ51bp). It is notable that these mutations evolved in PCA-adapted lineages under low-carbon conditions. This suggests that evolution in this environment may be limited by substrate uptake, potentially indicating diffusion or transport constraints. To understand how these mutations might affect the protein, we created an AlphaFold model of the porin (**Figure S4**). Our model showed that all deletions are located at the pore cap and, hypothetically, affect substrate transport.

### PHB accumulation hinders growth in aromatics

Given that the β-ketoapdipate pathway flows into the PHB cycle (**Figure 4A**), the discovery of recurrent mutations in *acsA*, the final enzyme in the PHB cycle (26), as well as mutations in putative regulators *ptsN* and *hprK,* collectively suggested that a metabolic imbalance of PHB may occur during growth on aromatics. To test this hypothesis, we characterized PHB accumulation for the mutants implicated in PHB cycling and for the ancestor. We measured PHB as a percentage of cell dry weight (CDW) in the exponential and stationary phases during growth on 5 mM VA.

Each of the evolved strains and the ancestor accumulated considerable amounts of PHB during the exponential phase (**Figure 4C**), suggesting a bottleneck somewhere along the PHB and EMC cycle. However, upon reaching stationary phase, all the evolved isolates with suspected PHB cycle mutations (*ptsN*, *acsA*, *hprK*) exhibited significantly lower (<1% of CDW) PHB content compared to the ancestor, which maintained ∼20% of CDW as PHB even in stationary phase. This is surprising, as 5 mM VA corresponds to a C:N ratio of 4:1, which is significantly below the threshold considered to be carbon excess conditions (C:N of ∼12:1) when PHB accumulation is typically activated (32). As a control to test the hypothesis that the PHB cycle affects the consumption of aromatics, we interrupted the PHB cycle by deleting *phaC*, the gene responsible for the polymerization of PHB (**Fig 4A**). This Δ*phaC* strain, as expected, showed no accumulation of PHB during both exponential and stationary phase.

We next tested whether reduced PHB accumulation alone can improve growth on aromatics. We observed that the beneficial mutations in *ptsN*, *hprK*, and *acsA* (genes involved in the PHB cycle) all contributed to increased growth rate on 5 mM VA (**Figure 4B**). Surprisingly, the *ΔphaC* ancestor strain, which lacked any evolved mutations, grew nearly as well as the evolved isolates. This suggests that PHB accumulation hinders growth on aromatics and that regulating the pathway (via mutations in *hprK*, *ptsN*) or increasing depolymerization (overexpression of *acsA*) enhances growth rate on aromatics.

Although not suspected of directly affecting the PHB cycle, during VA growth an isolate with the *epgR* mutation (from the PCA population) showed higher PHB accumulation in the exponential phase compared to the ancestor, but a reduction in PHB in the stationary phase. This suggests a different balance between PCA and VA for PHB accumulation, which may be related to the additional reducing power provided by the methoxy group and the NADH/NADPH balance in the PHB cycle. This *epgR* isolate also exhibited an improved growth rate on 5 mM VA.

To determine whether these mutations result in differential accumulation of PHB on non-aromatic carbon substrates, we tested PHB accumulation in these strains grown on succinate with a 100:1 C:N ratio (**Figure S6**). Here, carbon is present in ∼10 times excess, which should promote PHB accumulation in the stationary phase as nitrogen becomes limiting. Interestingly, the isolates followed a different pattern. While the *ptsN* and the *epgR* mutations still affected PHB accumulation, but only reduced accumulation by half compared to the ancestor, while isolates with mutations in *acsA* and *hprK* did not affect PHB accumulation (**Figure S6**). This suggests that specific regulatory pathways or enzymes selected for in evolution are condition specific.

### Evolution on aromatics reveals unexpected tradeoffs and asymmetric cross-adaptation

To assess the pleiotropic effects linked with adaptation to a novel pathway, we measured the growth rates of a subset of the evolved isolates on VA, PCA, methanol, and succinate. An initial hypothesis would be that evolution on VA would yield synergistic improvements on PCA, as VA metabolism encompasses all the steps involved in PCA catabolism, with the sole addition of a demethoxylation step that releases formaldehyde. Contrary to this expectation, all high VA evolved isolates exhibited growth defects at high concentrations of PCA, highlighting a surprising tradeoff, while in low PCA, there is no tradeoff for these mutations (**Figure 5, Figure S7**). Thus, we observed a concentration-dependent substrate-specific tradeoff, rather than the anticipated cross-benefit.

**Figure 5.**
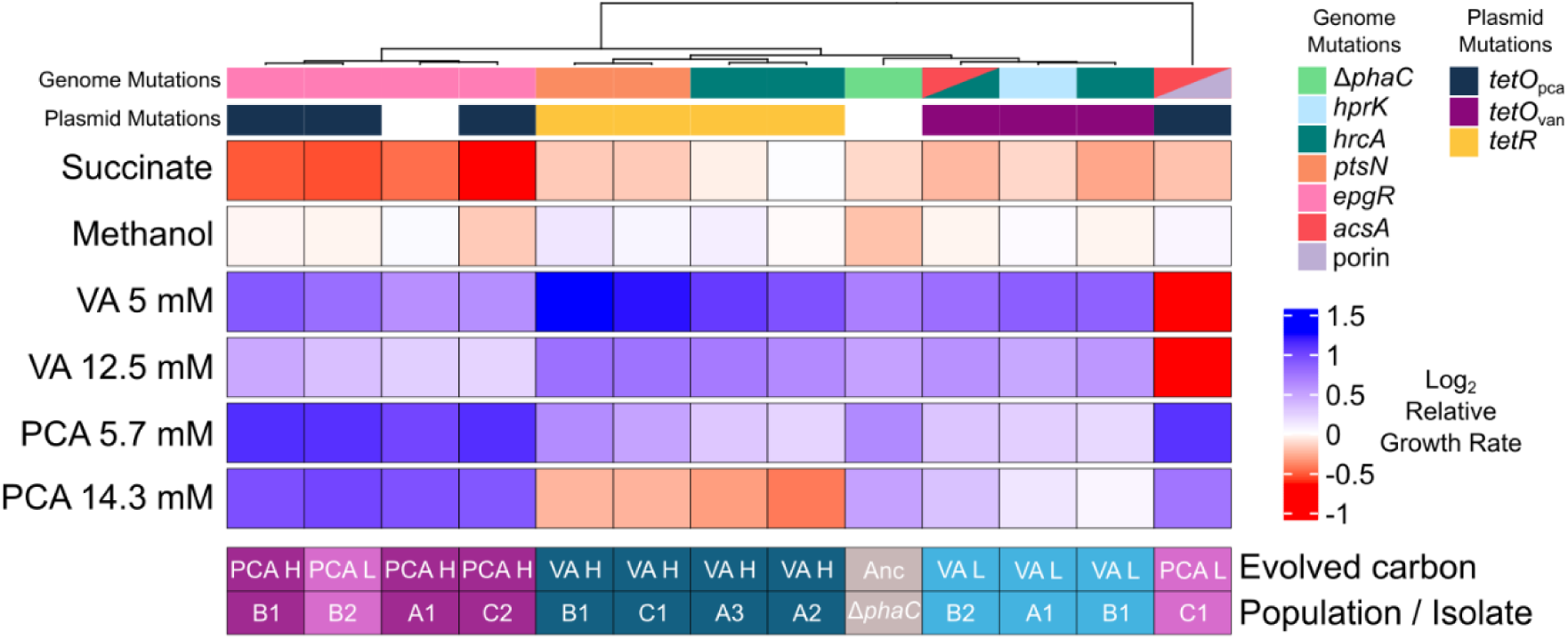
Relative growth rates across conditions. All isolates were grown in each evolved condition and a standard condition of succinate and methanol. Growth was relative to the ancestor growth rate in the corresponding condition and log_2_-transformed to better scale rates. Isolates (columns) were clustered using Euclidean distance measurements of the relative growth rates. Most isolates exhibit improved growth in all aromatic conditions compared to their ancestor. Trade-offs were seen in VA evolved strains that contained the *tetR* mutations (yellow bar) did worse in PCA and PCA evolved isolate with the porin mutations can no longer grow on VA. Isolates do either similarly or slightly worse on succinate compared to ancestor, while PCA evolved isolates with the *epgR* mutation (pink bar), do much worse. No significant growth differences were observed on methanol for any strain (See **Table S6** for TukeyHSD statistics).

As PCA is downstream of VA, evolution on PCA could lead to neutral or antagonistic effects, as demethoxylation is a key step in aromatic metabolism, or lead to synergistic pleiotropy, as increasing flux downstream could also help VA metabolism. Both scenarios appeared in the evolution of PCA, as most PCA evolved isolates exhibit improved growth on both PCA and VA (**Figure 5**). However, mutations within the outer membrane porin had the opposite effect, which completely lost the ability to grow on VA, regardless of the corresponding plasmid or other genome mutations (**Figure S5**).

While synergistic and antagonistic pleiotropy was seen between aromatic pathways, we were also interested in the effects of evolution on non-aromatic substrates. We observed that most evolved isolates exhibited lower growth rates on succinate than the ancestral strain **(Figure 5).** This was especially stark for PCA evolved strains with the *epgR* mutation, whose function is still a mystery; however, the gene neighborhood suggests central metabolism could be affected (**Figure 5 and S2B**). This tradeoff did not extend to methanol, however, where growth rates were largely unchanged for the evolved isolates.

Collectively, our results demonstrate that there are trade-offs within the “nested” pathway of the β-ketoadipate catabolism, where adaptation to one substrate may hinder the catabolism of another. Substrate-specific stressors, regulatory rewiring, and metabolic priorities shape these unexpected tradeoffs and cross-adaptations.

## Discussion

Experimental evolution of engineered *M. extorquens* on lignin-derived aromatics led to substantial improvements in growth within the selective environment, demonstrating the organism’s capacity for rapid adaptation to novel and challenging substrates. As shown before, this result confirms that integration of even a functional foreign pathway benefits significantly from evolutionary fine-tuning (17, 33), especially under substrate-specific pressures.

The adaptive mutations in the TetR/*tetO* system demonstrate how the flexibility of operon regulation can be a key evolutionary driver under substrate-specific selection. Rather than increased expression of both gene cassettes being universally beneficial, the system evolved different expression strategies tailored to the demands of each substrate (**Figure 3**). This highlights two salient points: first, catabolic flux through the introduced pathway is only part of the equation, and it must be matched to the cell’s tolerance for expression burden; and second, the context of enzyme complexity shapes whether global (TetR) or operon-specific (*tetO*) changes are optimal.

For evolution on aromatics, the regulatory specificity was most likely due to the burden of VanAB, the multi-component demethylase complex required for VA conversion, which has proven challenging to express and purify in heterologous systems, with low yields of active protein even under optimized conditions (34). Such protein-folding stress likely explains why several VA-evolved isolates acquired mutations in *hrcA*, a transcriptional repressor of heat shock chaperones like GroEL and DnaK (30, 35–37). While direct derepression of these chaperone systems was not seen, other systems could buffer proteostatic stress associated with elevated *vanAB* expression.

Interestingly, PCA-evolved strains also performed well on VA, despite lacking direct mutations in the *van* operon or other regulatory elements. A likely explanation for the cross-adaptation of PCA-evolved strains to VA lies in the near-universal presence of mutations in *epgR*. These mutations may have optimized carbon flux through the TCA cycle, thereby improving the cell’s ability to process intermediates derived from either PCA or VA. Therefore, *epgR* mutations may have enhanced downstream metabolic capacity, allowing PCA-evolved strains to better handle VA catabolism even without direct changes in *van* operon regulation. These findings reinforce the idea that global cellular adaptations, particularly those that improve downstream flux handling, can offset the lack of pathway-specific optimization (17, 38).

Despite improved growth on VA and PCA, most evolved isolates showed diminished growth on succinate compared to the ancestral strain. This reveals a specialization-versus-versatility tradeoff, in which adaptive mutations that enhanced aromatic catabolism appear to have reconfigured central metabolism in ways that impaired the use of a generalist substrate. This tradeoff was particularly striking among PCA-evolved strains, all of which carried mutations in *epgR*. If a bottleneck exists in central metabolism, increasing flux through these pathways would increase growth on the new substrate, but might lead to an imbalance on already fast substrates such as succinate.

Perhaps one of the greatest insights from the evolved isolates was the novel connection between aromatic degradation and PHB synthesis. Mutations in the transcriptional start sites of *acsA*, found in both VA and PCA strains, point to a shared pressure to modulate acetoacetate metabolism. VA-specific mutations also affected C:N regulatory systems, reinforcing the idea that aromatic catabolism rewires cellular priorities, redirecting resources away from storage (PHB) and toward biomass accumulation. It was surprising that the ancestral *M. extorquens* strain stored substantial amounts of PHB (up to 20% of cell dry weight) during growth on 5 mM VA, despite this condition corresponding to a low C:N ratio (∼4:1), well below the threshold typically associated with carbon excess (32). This suggests that PHB synthesis was not triggered by surplus carbon, per se, but instead reflects overflow metabolism, a cellular strategy to divert excess reducing equivalents or acetyl-CoA into storage when central metabolic flux is constrained (39, 40). In this case, redox imbalance or bottlenecks introduced by aromatic degradation may have prompted PHB accumulation as a protective sink (41). This phenomenon might be unique to *Methylobacterium* due to the Kce enzyme, which produces acetoacetate and succinyl-CoA at the end of the β-ketoadipate pathway linking aromatic degradation directly to PHB storage metabolism rather than the TCA cycle. We see that evolution selected for increased growth by restructuring central metabolism, as evolved isolates stored less than 1% PHB at stationary phase. These strains matched or exceeded the growth performance of a Δ*phaC* mutant, which entirely lacks PHB synthesis. This metabolic shift likely liberated NADPH, acetyl-CoA, and other critical intermediates, increasing carbon and energy availability for core functions. However, we know that PHB supports survival in the environment, especially during starvation (42, 43), suggesting another potential trade-off in adaptation to aromatic metabolism.

While the acquisition of a foreign metabolic pathway via horizontal gene transfer can unlock the ability to exploit a novel substrate, this benefit often comes at the expense of ancestral functions, leading to antagonistic pleiotropy. We demonstrate this here by showing the complex trade-offs involved in integrating aromatic metabolism with central metabolism in a model methylotroph. As evolving populations optimize growth on the new substrate, these trade-offs introduce dynamic shifts in the fitness landscape, leading to incremental mutations yielding disproportionately large changes in resource utilization profiles. Over successive rounds of selection, the cumulative burden of such trade-offs drives increasingly pronounced phenotypic and genotypic divergence from the ancestor (44). Ultimately, this process of niche specialization magnifies lineage differentiation, leading to speciation, and illustrates how the evolution of foreign pathways embeds tradeoffs that can propel microbial diversification.

## Supplemental Figures

**Figure S1.**
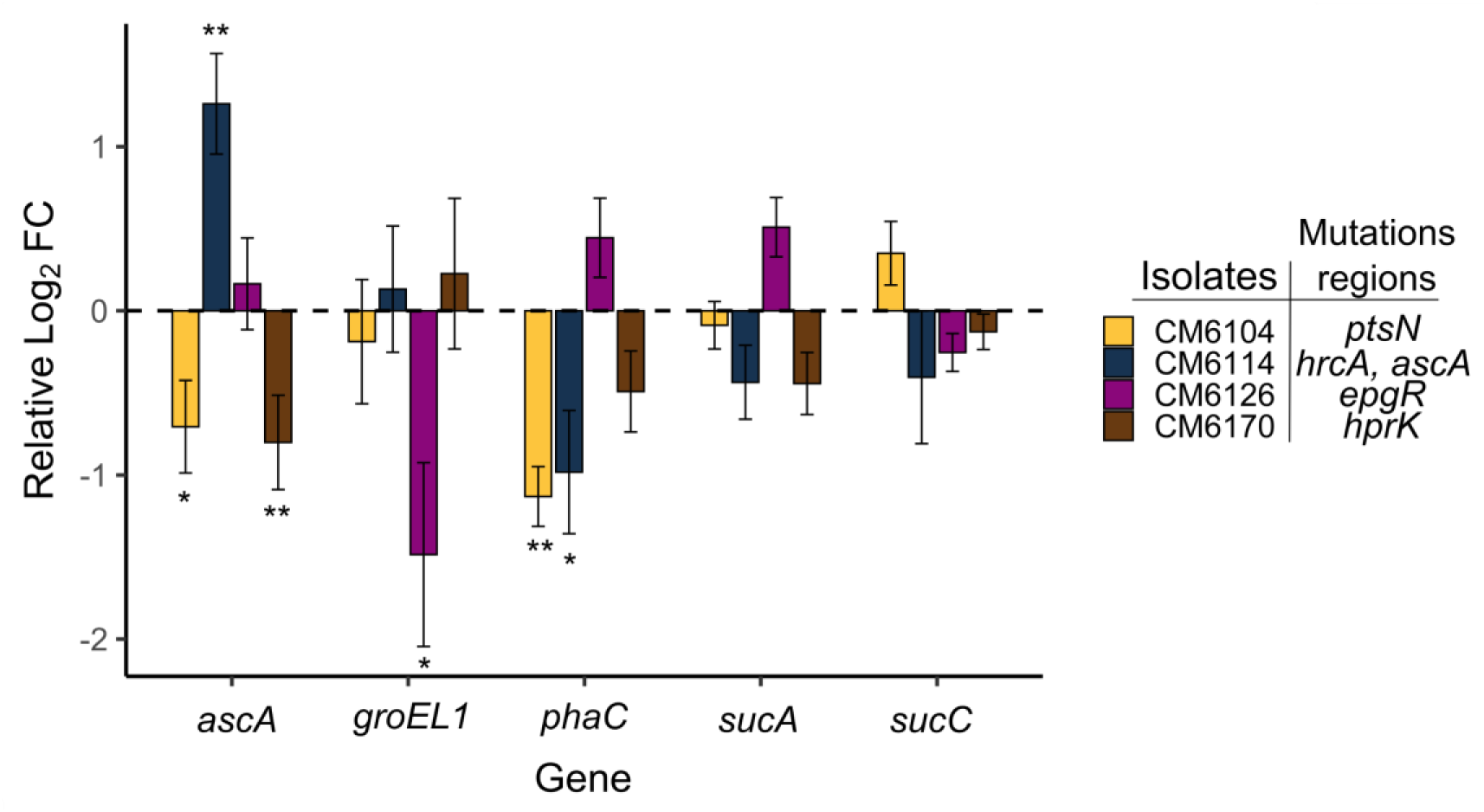
Relative expression of genes regulating central metabolism compared to the ancestor grown on VA. Genes *ascA* (3-oxoacid transferase/acetoacetate-CoA transferase), *phaC* (poly-3-hydroxybyturate polymerase) are involved in the PHB cycle. Mutations in the regulation of *ascA* (CM6114, −120 bp from start codon) significantly upregulate the gene, while putative PHB cycle regulator mutations CM6170 (*hprK*_S133G) and CM6104 (*ptsN*_(C)_5->4_) downregulate *ascA*. While *phaC* is downregulated in all three PHB cycle mutations, but is only significant with *ptsN* and *ascA* mutations. Isolate CM6114 carried the *hrcA* mutation (Δ96 bp), which we suggested regulated the *groEL* complex; however, we did not observe an effect in this isolate, whereas we did observe a downregulation in isolate CM6126 (*epgR*). We proposed that the *epgR* gene, which is a *tetR*-family regulator sitting in front of many TCA cycle genes, may regulate some of the intermediate enzymes of this cycle. However, both *sucA* (oxoglutarate dehydrogenase) and *sucC* (succinate-coA ligase) do not show any significant changes between strains. Bars represent mean relative log₂ fold change ± standard error (SE) from three biological replicates. Asterisks (*) indicate statistically significant differences in gene expression relative to the ancestor (* p < 0.05, ** p < 0.01, *** p < 0.001, **** p < 0.0001, two-tailed t-test).

**Figure S2.**
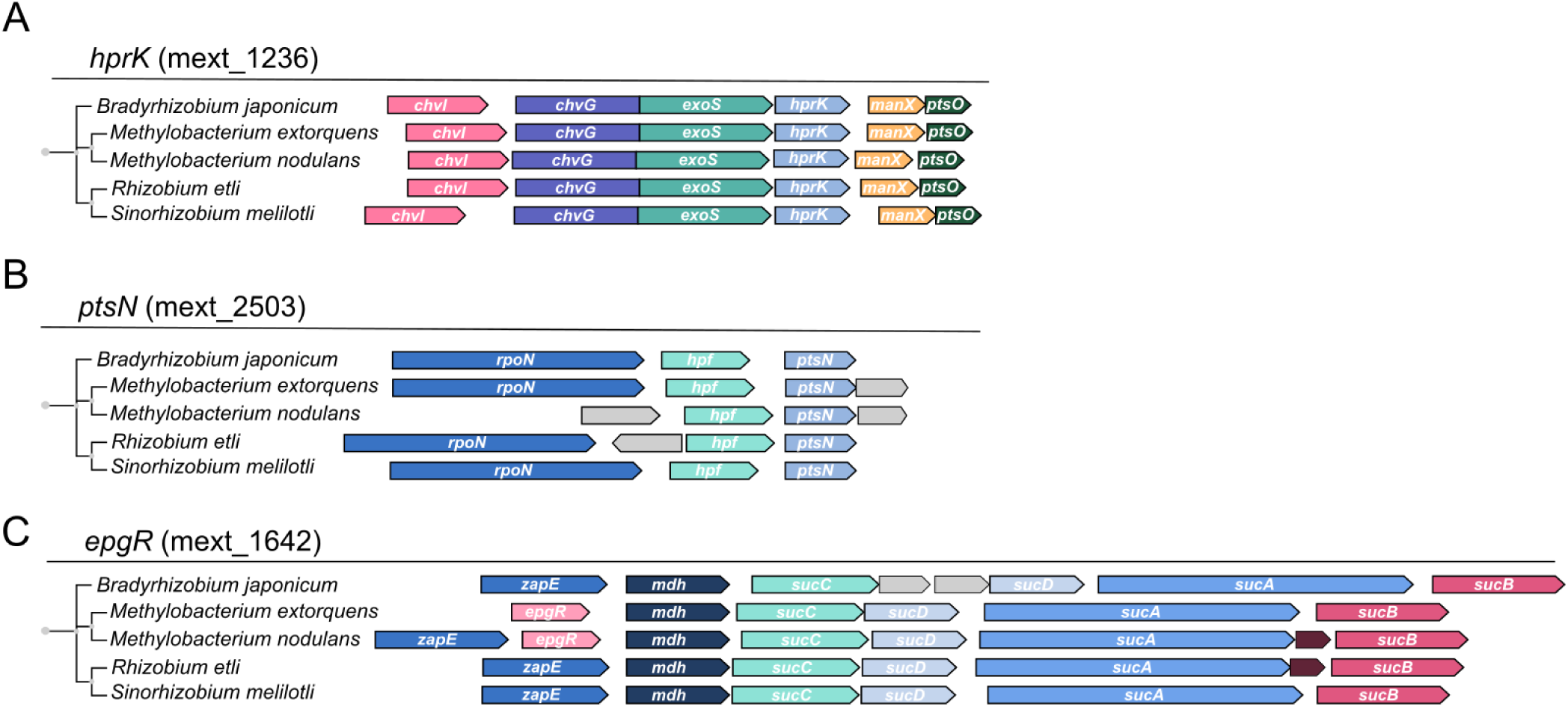
Alignment of genes selected for in the aromatic evolution experiment compared to other well-studied Hyphomicrobiales. A) HPR kinase (hprK, *mext_1236*) is a part of the PTS signal cascade, which has been shown to regulate PHB production in other Hypomicrobiales. The PTS cascade consists of a membrane protein (*chvG*/*exoS*) and a series of kinases (*chvI*, *hprK*, *manX*/*ptsO*). B) The gene *ptsN* is also potentially involved with the PTS cascade, but it is not associated with the other components of the cascade. C) The novel *epgR* (Enhanced PCA growth regulator) gene was found to be unique to the *Methylobacterium* genus and sits directly in front of multiple TCA cycle genes.

**Figure S3.**
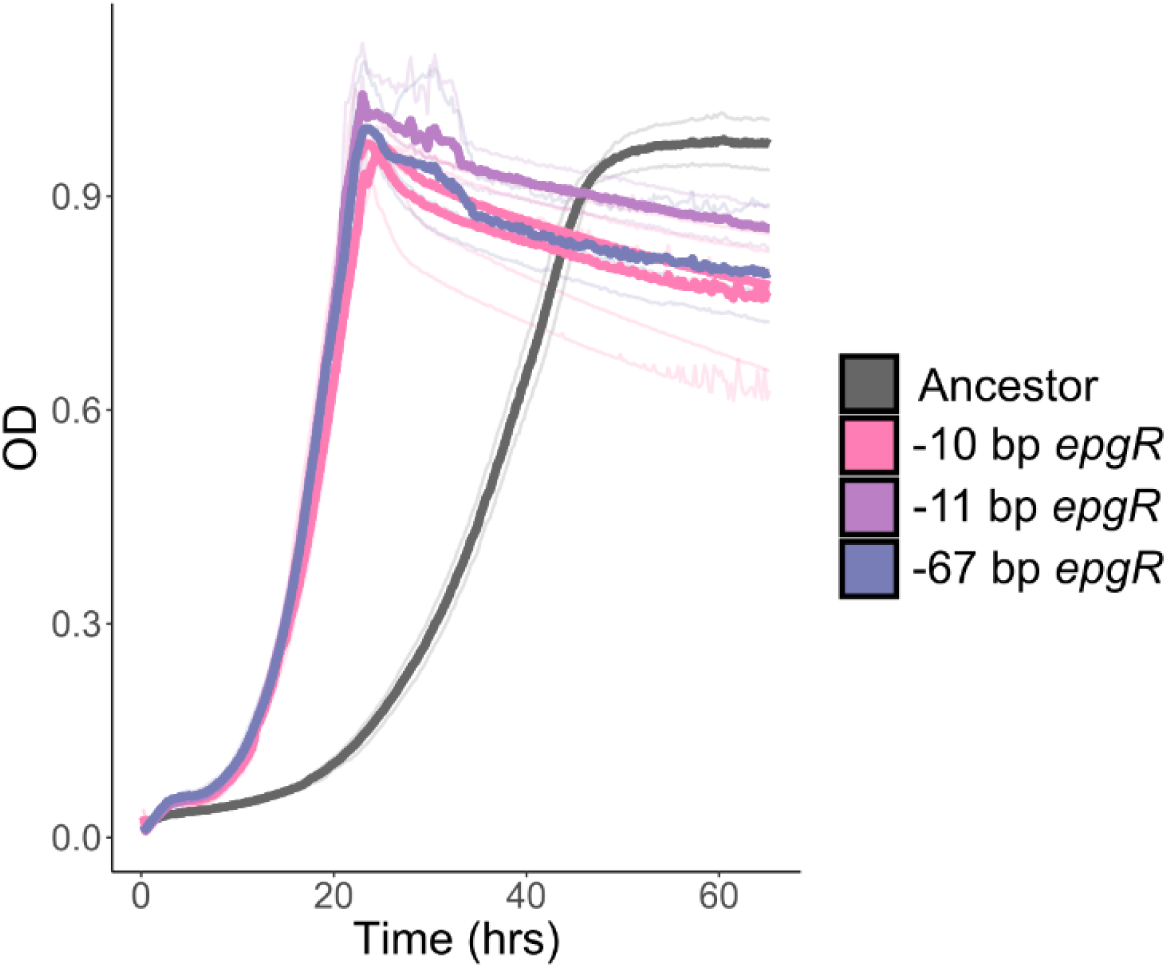
Growth of PCA evolved isolates with *epgR* mutations on high PCA conditions. Mutations in the transcription of *epgR*, either in the −10, −11, or −67 bp position in the gene, are equivalent for growth on PCA.

**Figure S4.**
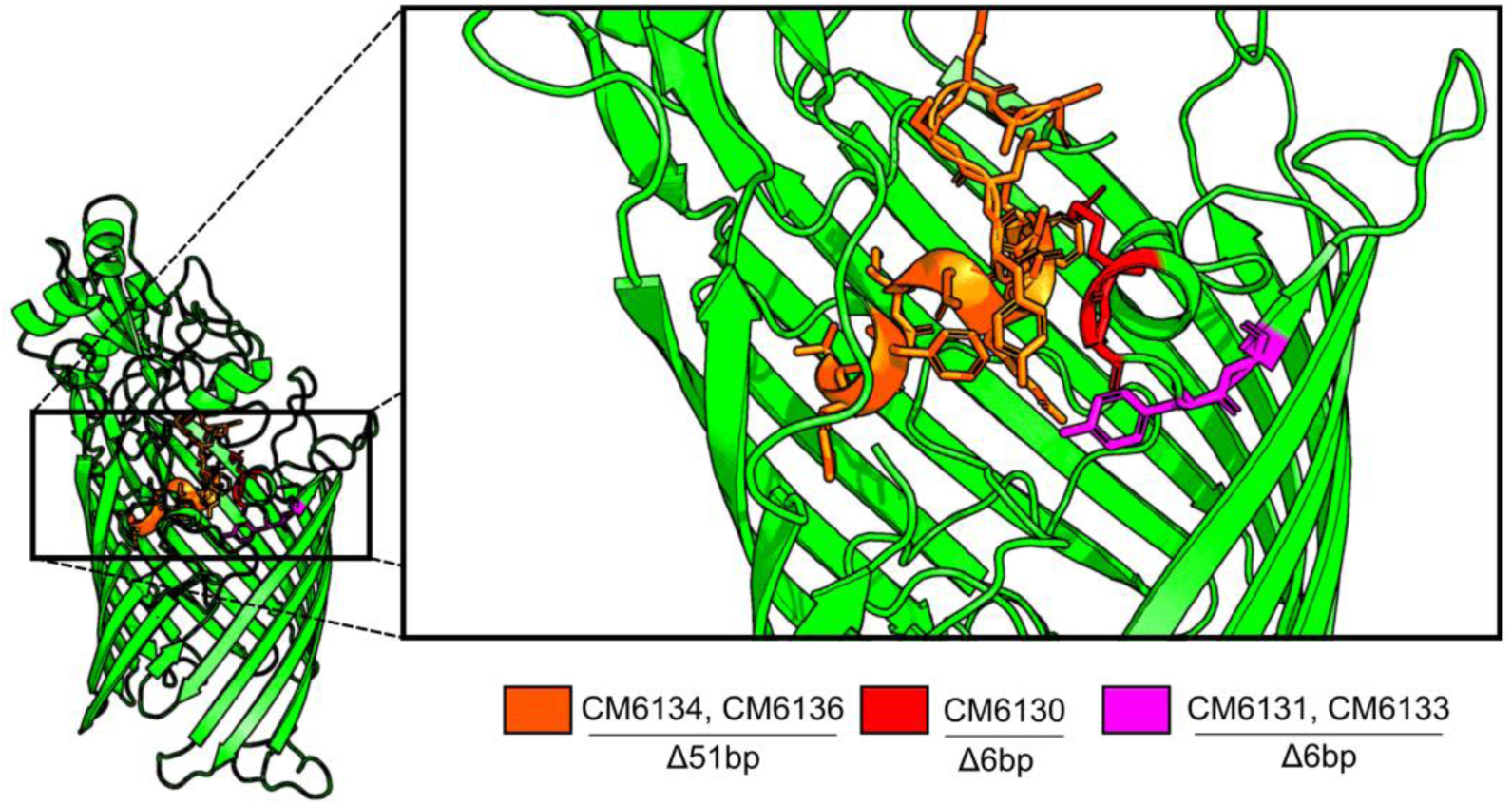
Alpha Fold structure of porin and deletions found in strains from the evolution experiment. In-frame deletion of the porin gene leads to the removal of multiple amino acids, all located at the top of the β-barrel structure. No information is known on the substrate that is transported, but removal of any of these “caps” of the porin is shown to be detrimental to VA growth.

**Figure S5.**
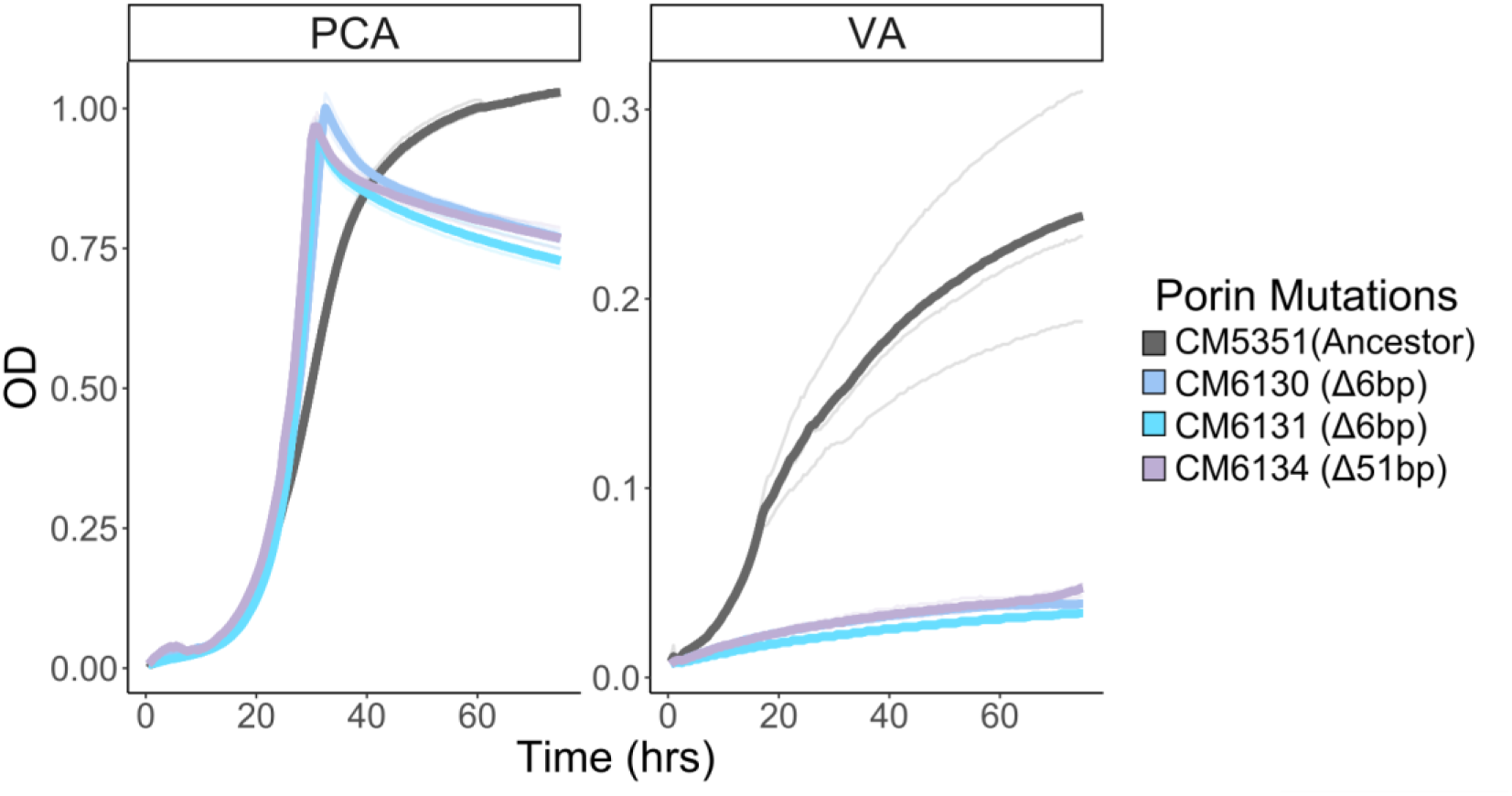
Growth of porin mutants on PCA and VA. The porin in-frame deletions lead to a removal of the “cap” on top of the β-barrel. This leads to an increased growth rate on PCA, while it is detrimental to the growth on VA.

**Figure S6:**
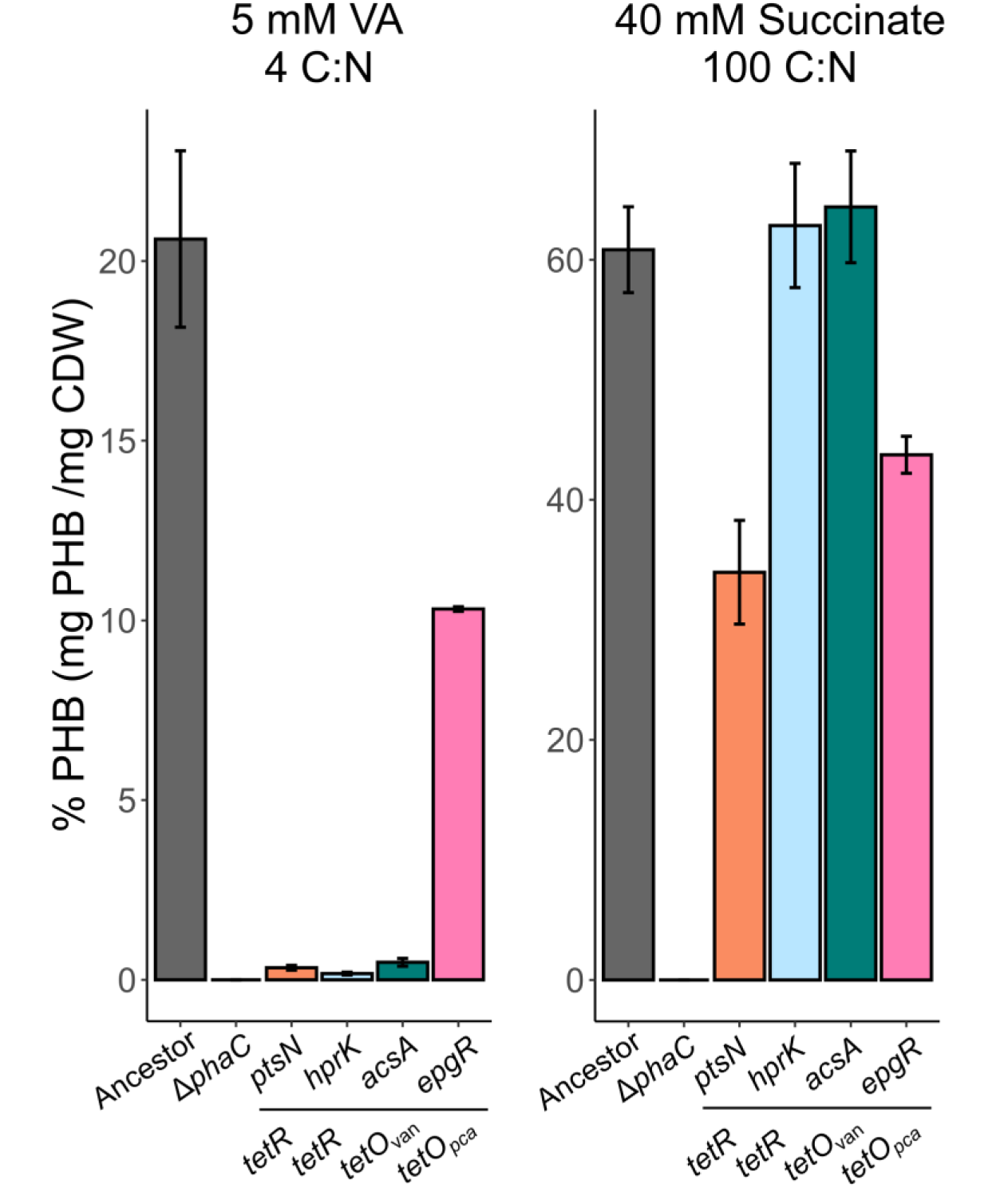
PHB accumulation on high C:N ratio of succinate during the stationary phase. The ancestor and *epgR* mutants both accumulate PHB in stationary phase when grown on 5 mM VA, which is at a 4 C:N ratio (left). When grown at a 100 C:N ratio, we observe that *pstN* and *epgR* mutations decrease PHB production in succinate, whereas *hprK* and *acsA* have no effect (right). This shows substrate-specific adaptation to aromatics rather than a general PHB regulation.

**Figure S7.**
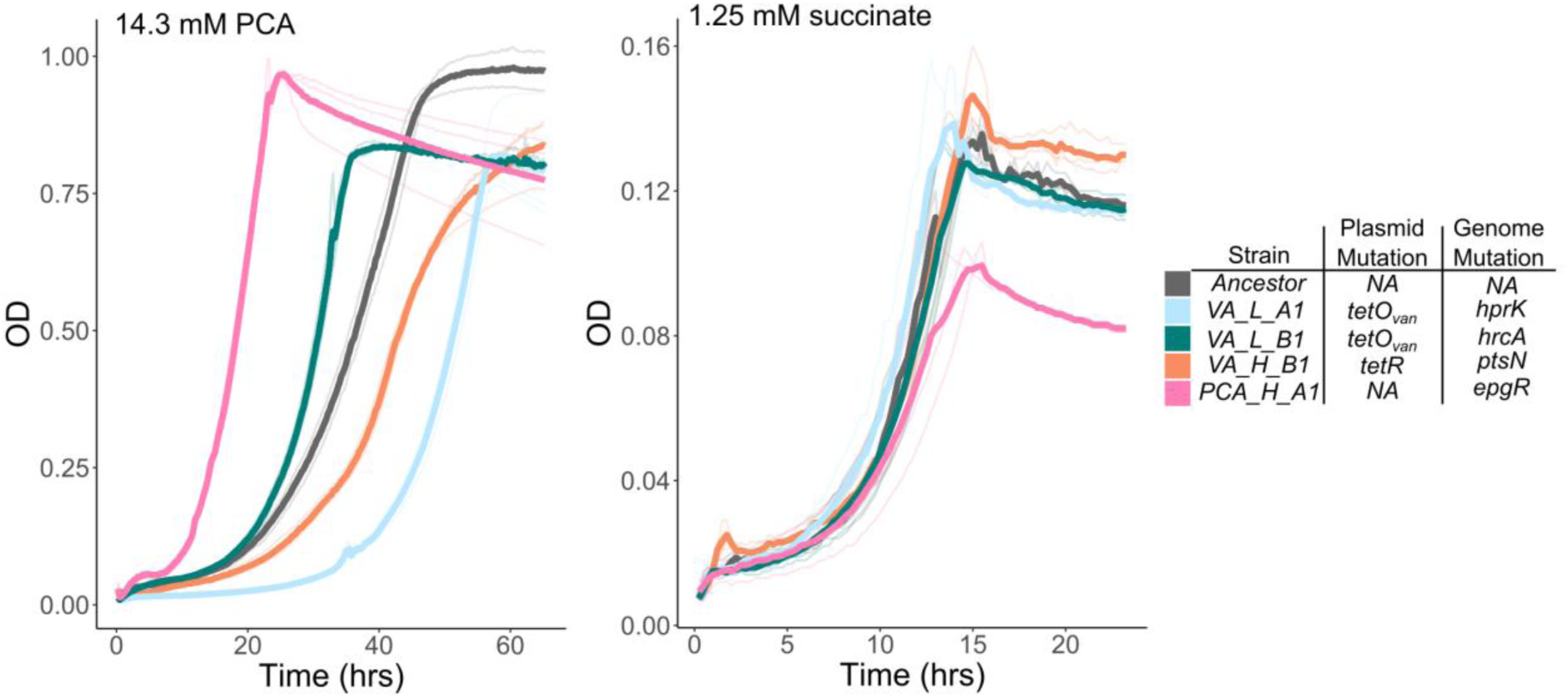
Growth of VA and PCA evolved isolates on 14.3 mM (high) PCA and 1.25 mM succinate. A) VA isolates show impaired growth on high PCA, relative to the ancestor, indicating a tradeoff. Some of which contain *tetR* mutations in the plasmid. B) The evolved strains grow poorly on succinate, especially the high PCA evolved strains, which have the *epgR* mutations.

## Supplemental information

**A) CRICE element (red) present in both *groES* operons in *Methylobacterium extorquens* PA1**

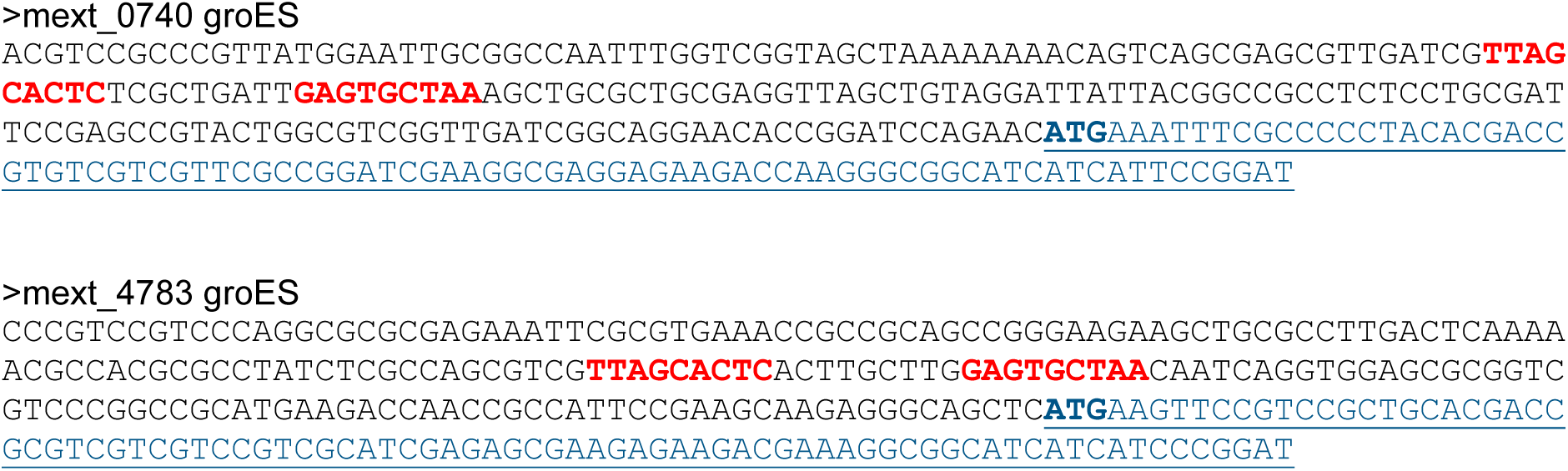

**B) Point mutations (C->T, red) in front on the *tetR*-like regulator *ameR* (now called *epgR*)**

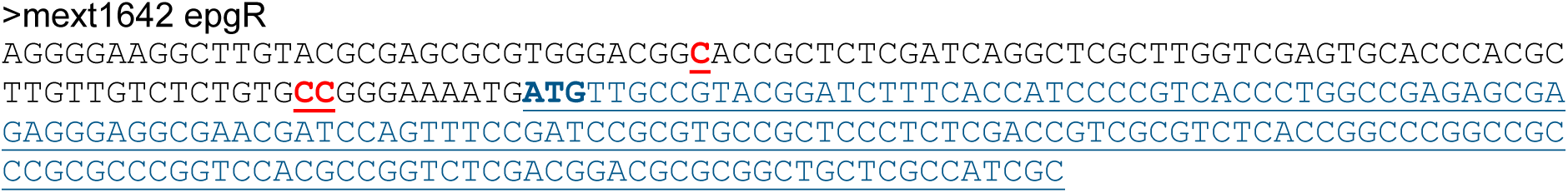

